# Efficacy of natural marine sponges as a passive environmental DNA sampler for freshwater fish diversity monitoring

**DOI:** 10.1101/2025.06.05.658201

**Authors:** Ryohei Nakao, Manami Inaba, Seiji Miyazono, Minoru Saito, Keita Maruyama, Fumiko Imamura, Yoshihisa Akamatsu

**Affiliations:** Graduate School of Science and Technology for Innovation, Yamaguchi University, Yamaguchi University, 2-16-1, Tokiwadai, Ube, Yamaguchi, Japan; Japan International Research Center for Agricultural Sciences, 1-1 Ohwashi, Tsukuba, Ibaraki, Japan; Research and Development Center, Nippon Koei Co., Ltd, 2304, Inarihara, Tsukuba, Ibaraki, Japan; Department of Ocean Sciences, Tokyo University of Marine Science and Technology, 4-5-7, Konan, Minato-ku, Tokyo 108-8477, Japan

**Keywords:** environmental DNA, metabarcoding, passive sampler, river ecosystem, freshwater fish

## Abstract

Environmental DNA (eDNA) analysis is a cost-effective and noninvasive tool for species and biodiversity monitoring in aquatic environments. Passive eDNA sampling is a novel alternative to conventional sampling methods such as water filtration. In this study, we examined the efficacy of the sponge skeleton as a passive eDNA sampler for monitoring freshwater fish. The performance of the passive sampling method was compared with that of standard water filtration in a river environment. Five DNA extraction methods were used in three experiments, and a suitable method for DNA extraction from sponge skeleton was identified. Quantitative fish metabarcoding using MiFish primers revealed no significant differences in species richness between the aqueous and passive sampling methods. Although both the sampling methods showed comparable trends in fish community structure, different clusters were identified for water sampling and passive samplers based on the differences in the DNA concentration of each fish species. The fish diversity in the passive samples was comparable among four DNA extraction methods (except for the filtration method) using the direct capture method. Our results demonstrate the efficacy of passive eDNA sampling for monitoring freshwater fish diversity and the potential use of sponge skeletons as absorption materials for passive eDNA samplers.

## Introduction

Environmental DNA (eDNA) analysis is increasingly being used to monitor biodiversity in various ecosystems. This method can detect diverse organisms by capturing and extracting DNA from excrement and extracellular DNA from environmental samples such as water, soil, and air (Turner et al. 2015, Tsuji et al. 2019, Lynggaard et al. 2022). Environmental DNA analysis is a sensitive, non-invasive, and broadly applicable tool for species detection. Environmental DNA metabarcoding can simultaneously detect multiple species (Miya et al., 2015; Deiner et al., 2017). It is effective for performing rapid and large-scale multispecies surveys (Olds et al., 2016; Yamamoto et al., 2017). In particular, eDNA metabarcoding has been used to monitor the fish diversity in various water bodies ranging in size and type of aquatic ecosystems (Hänfling et al. 2016, Nakagawa et al. 2018, Oka et al. 2021). Many studies have shown that eDNA metabarcoding is a more effective and sensitive method for fish diversity assessment than conventional approaches such as visual census, fyke net capturing, and electrofishing (Yamamoto et al. 2017, Shaw et al. 2016).

Water eDNA is an environmental DNA collected from water samples. It is most commonly used to monitor fish diversity. Water eDNA from fish is generally collected and concentrated from water samples by filtration through artificial membranes (Yamamoto et al. 2017, Ushio 2019, Jeunen et al. 2019). However, the fish diversity in water eDNA is generally snapshot information. Moreover, insufficient sampling efforts could result in an underestimation of the fish diversity (Zinger et al. 2019). For example, if the concentration of the target DNA is low (such as for rare species), small sampling volumes or few replicates can yield false negatives. Furthermore, because eDNA concentrations reflect the activity levels of the target species (e.g., diurnal or nocturnal), the mismatch of timing between water sampling and fish ecology may increase false-negative detections. Large volumes of water filtration and sampling replicates are required to remove false negatives and provide reproducibility of fish diversity. However, the conventional sampling method using water filtration requires considerable time and cost to achieve such improvements.

Passive eDNA collection methods directly submerge the sampling material in a water body. These facilitate reproducible biomonitoring as a simpler and more cost-effective sampling method than conventional active filtration methods (Bessey et al., 2021; Kirtane et al., 2020; Zhang et al. 2024). The absorption materials for passive eDNA sampling used in previous studies generally have porous substances, sufficient adsorption capabilities to detect the target eDNA, and simple or user-friendly forms. The submersion materials used in passive eDNA sampling methods include activated carbon, montmorillonite (Kirtane et al., 2020), living sponges (Mariani et al., 2019; Cai et al., 2022), and biofilms (Rivera et al., 2021). In the case of artificial materials, Verdier et al. (2022) created the passive, 3D-printed, and convenient-to-use eDNA sampler using hydroxyapatite (a natural mineral with a high DNA adsorption capacity). Yan et al. (2024) deployed passive sampling using GF filters by the angling (short-term immersion at the site) and trolling (towing in the water) methods in large lake environments. These studies showed that the effectiveness of passive collection methods in detecting fish is similar to that of conventional active filtration methods.

Furthermore, these studies provided similar estimates of total fish biodiversity. In addition, because passive eDNA samplers require minimal or no supporting equipment (e.g., no vacuum or peristaltic pumps), these are suitable for deployment in remote environments and by non-experts.

Natural and artificial sponges are likely to be superior passive eDNA samplers owing to their universal distribution, high regeneration capability, and convenience of sampling. For example, filter-feeding organisms have been shown to naturally accumulate eDNA in their tissue matrices (Jeunen et al., 2021). Mariani et al. (2019) showed that eDNA extracted from live marine sponges could detect various marine fishes and mammals in the Mediterranean Sea and Antarctic Ocean. Turon et al. (2020) highlighted the suitability and cost-effectiveness of live marine sponges as passive eDNA samplers for describing tropical fish communities from coral reefs in Southeast Asia. Jeunen et al. (2022) indicated that more porous passive filter materials such as artificial sponges could potentially achieve higher detection rates through the adsorption and retention of less abundant DNA and DNA bound to particulate matter. Therefore, natural sponge skeletons could potentially be used as effective materials for passive eDNA samplers. However, the knowledge of the application of sponge skeletons to eDNA studies is insufficient for both marine environments and freshwater environments, particularly in rivers.

In this study, we examined the efficacy of the sponge skeleton as a passive eDNA sampler (sponge sampler) in freshwater river environments. We submerged sponge samplers in a river and estimated the freshwater fish diversity from the sponge eDNA samples using eDNA metabarcoding. In addition, to examine the performance of eDNA accumulation with long-term immersion of sponges, we also collected river water eDNA samples (the major material in eDNA surveys) in parallel with sponge eDNA samples and compared the differences in the number of fish species and fish species composition between passive and water samples. Furthermore, several sampling and DNA purification methods for passive samplers were examined to estimate the optimal passive sampling method using sponge skeletons.

## Materials and Methods

### Study field and environmental DNA samplings

In this study, field surveys were conducted in the middle reaches of the Saba River Basin, Yamaguchi Prefecture, Japan, on July 30–31, October 3–4, 2021, and May 9–10, 2022 (Fig. 1A, B). The fish activity times in rivers differ across species (e.g., diurnal or nocturnal). Therefore, the species composition detected may vary depending on the timing of water sampling. Sponge-passive samplers can potentially overcome these variabilities in species detection through long-term eDNA adsorption onto the sponge skeleton in water. To accurately assess the function of eDNA adsorption on the passive sampler, we conducted water sampling multiple times during the immersion of the passive sampler. For water sampling, 3 × 1 L of surface water was collected directly from the center of the riffle using bleached plastic bottles every 2 h (0, 2, 4, 6, 8, 10, 12, 14, 16, 18, 20, 22, and 24 h in 2021) or 3 h (0, 3, 6, 9, 12, 15, 18, 21, and 24 h in 2022) after submerging the equipment (Fig. 1C). In this study, all the experiments were started at 11.00. Therefore, in Experiments 1 and 2, we defined 0–6 h (11:00–17:00) and 20–24 h (7:00–11:00) as daytime, and 8–18 h (19:00–5:00) as nighttime. In Experiment 3, we also defined 0–6 h (11:00–17:00) and 21–24 h (8:00–11:00) as daytime and 9–18 h (20:00–5:00) as nighttime. Water sampling was also performed downstream of the equipment to prevent contamination from riverbed disturbances caused by human movement.

**Fig 1.**
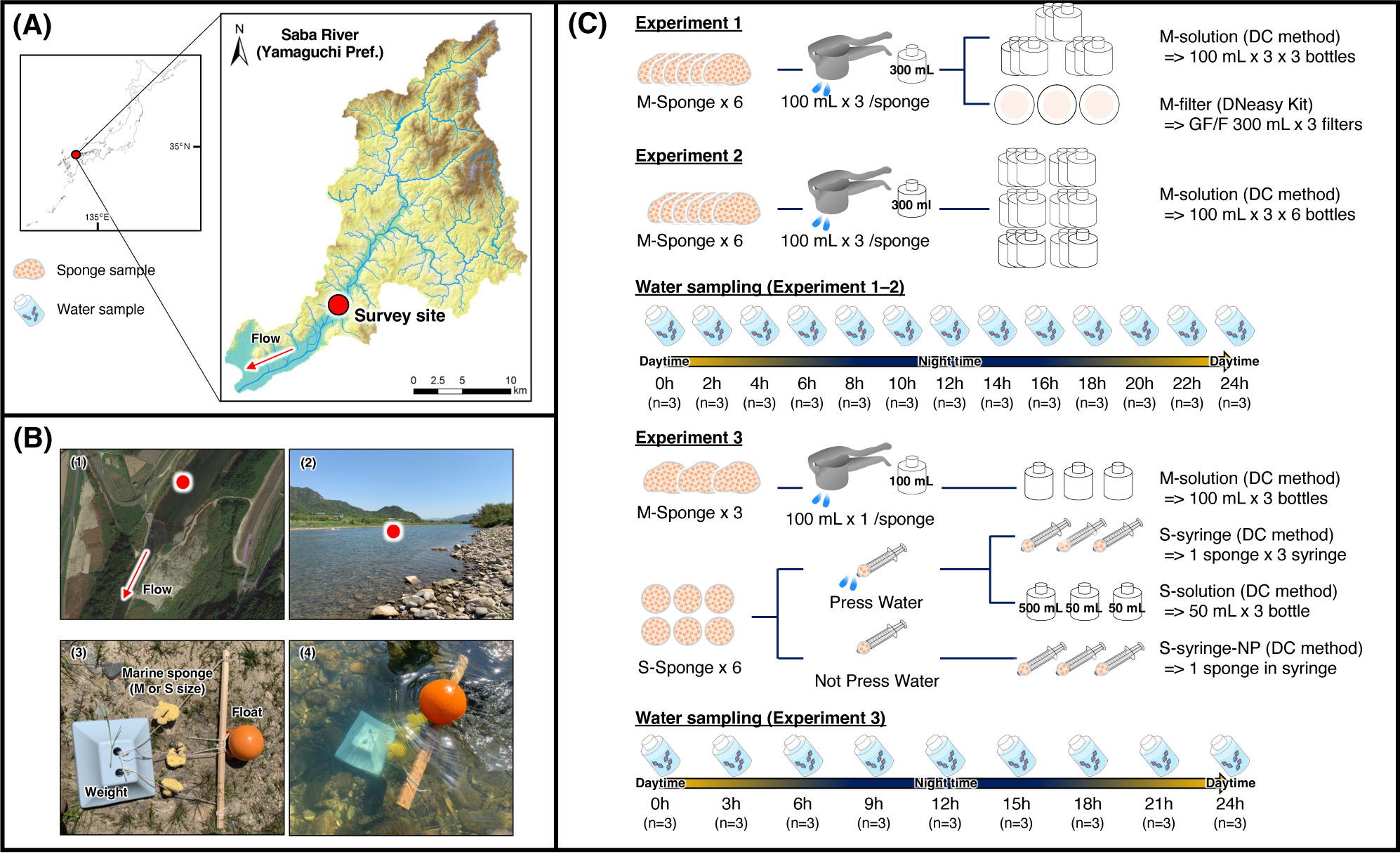
Study site information and descriptions of experimental design in this study. (A) Descriptions of survey site, (B) detailed information of survey site and passive sampler, (C) methodological procedures of water and passive sampling in each experiment

Water samples were filtered immediately on site using a portable pump (Sentino microbiology pump; Cytiva, Tokyo, Japan) and GF/F glass fiber filter (pore size 0.7 µm; Cytiva). After filtration, the equipment was bleached to prevent DNA cross-contamination between water samples. The filters were stored at −16°C in the ice box and taken back to the laboratory at 12 and 24 h, respectively. The filters were stored at −20°C until DNA extraction.

Specific passive sampling equipment was used to submerge the sponge samplers (Fig. 1B, C). This equipment consists of a weight and float connected by ropes. The sponge skeleton of the *Spongia* spp. species was used as the adsorption material for the sponge sampler. Sponge samplers were attached to the midpoint of the rope-tip wraps before submerging the equipment to enable the sponges to float in the middle layer of the water. The equipment was placed near the river thalweg for 24 h for passive eDNA sampling. After 24 h, the sponge samplers were collected from the equipment and stored according to specific procedures in each experiment. After the sponge samplers were collected, the equipment was removed from the river. In this study, two sponge sizes were used for passive samplers: medium (M-sponge; approximately 100 mm in diameter) and small (S-sponge; approximately 3–40 mm in diameter).

In this study, we compared various methods to determine the optimal method for recovering eDNA from sponge samples. In Experiment 1 (July 30–31, 2021), six M-sponge samplers were used, and 100 mL of the pressed solutions were extracted from each sampler using a handheld juicer (Fig. 1C). Next, 100 mL of distilled water was added to the M-sponge to obtain 100 mL of pressed solution. Distilled water was added, and re-extraction was performed two times. A total of 300 mL of the pressing solution was obtained. Three of the six pressed solutions were additionally separated into 3 × 100 mL aliquots using 100 mL polyethylene bottles (M-solution).

M-solutions were stored at 4°C in a cooler box and transferred back to the laboratory. Then these were stored at −20°C until DNA extraction. The 300 mL of the other three M-sponge solutions in Experiment 1 were filtered with GF/F. The filters were stored at −16°C in the ice box and transferred back to the laboratory. Then, these were stored at −20°C until DNA extraction (M-filter). Six M-sponge samplers were used in Experiment 2 (October 3–4, 2021). All the pressed solutions were handled as M-solution samples and stored using the methods in Experiment 1 (Fig. 1C). In Experiment 3 (May 21–22, 2022), four passive-sampling methods were used. Three M-sponge and six S-sponge samplers were submerged in the river and collected after 24 h. In Experiment 3, 100 mL of the first pressed solution was extracted from each M-sponge sampler and used as the M-solution 2 samples (Fig. 1C). Six S-sponge samples were collected from the equipment and stored in separate syringes. Three of the six S-sponge samplers were pressed into syringes and separated into syringe samples (S-syringes) and pressed solutions (S-solutions). Approximately 50 mL of the S-solution sample was extracted from each S-sponge sampler and collected in 100 mL polyethylene bottles. The remaining three S-sponge samplers were stored in unpressed syringes (S-syringe NP). In Experiment 3, all the passive samples were stored at 4°C in a cooler box and transferred back to the laboratory. Then, these were stored at −20°C until DNA extraction.

### DNA extraction from water and sponge samples

The total eDNA of the filter samples was extracted and purified as described by Tsuji et al. (2022). Briefly, each filter was placed in the upper part of a Salivette tube (Sarstedt, Nümbrecht, Germany) and pre-centrifuged at 5000 × g for 1 min to remove excess water. Each filter was soaked with 440 µL of a mixture composed of 200 µL of Buffer AL, 200 µL of H2O, and 20 µL of proteinase K, and incubated at 56°C for 45 min. Subsequently, the cartridges were centrifuged at 5000 × g for 3 min, and the extract solutions of each tube were collected. The total eDNA was purified from extract solution using DNeasy Blood & Tissue Kit (QIAGEN, Hilden, Germany) and eluted in 100 μL of Buffer AE from a spin column according to the manufacturer’s instructions.

The total eDNA of the solution samples in the passive sampling methods (M-solution and S-solution) was extracted and purified using Maxwell^®^ RSC Enviro Total Nucleic Acid Kit (Promega, Madison, WI, USA) according to the manufacturer’s instructions (hereafter, the direct capture (DC) method). Briefly, 12.5 µL of proteinase solution from the DC kit was added per 1 mL of pressed solution. The samples were incubated for 30 min at ambient temperature. The samples were then centrifuged at 3000 × g for 10 min at room temperature to remove the precipitate. The supernatant was divided equally into two 50 mL tubes. Furthermore, 6.0 mL of Binding Buffer 1 and 0.5 mL of Binding Buffer 2 were added to the supernatant. Then, 24 mL of isopropanol was added to the mixture, mixed gently, and passed through a PureYield Midi Binding Column using a VacMan Vacuum Manifold. Nucleic acids captured on the PureYield Midi Binding Column were washed with 5 mL of Column Wash 1, followed by 20 mL of Column Wash 2. Nucleic acid was eluted with 750 µL of pre-warmed (60°C) nuclease-free water. Then, the eluate was transferred to a Maxwell^®^ RSC Cartridge, and the cartridge was held in a Maxwell^®^ deck tray. DNA purification and elution of nuclease-free water were performed on a Maxwell^®^ instrument according to the manufacturer’s instructions. After the procedures were completed, the eluate was collected from the deck tray, and 100 µL of eDNA solution was obtained. The instrument was then subjected to UV light treatment to prevent cross-contamination.

The total eDNA of syringe samples in the passive sampling methods (S-syringe and S-syringe-NP) was extracted and purified using the extraction kit for the sponge solution samples with a marginal modification in the incubation step. Briefly, proteinase K powder (approximately 30 U/mg; Nacalai Tesque, Kyoto, Japan) was dissolved in TE buffer (pH 8.0) to prepare a proteinase K stock solution (10 U/mL). The DNA elution mixture was prepared by adding 2.0 mL Proteinase K stock solution to 18 mL TE buffer (pH 8.0). An S-syringe sample was submerged in this solution in each syringe and incubated for 45 min at 56°C. The solution sample was discharged to a 50 mL tube from each syringe. Then, 6.0 mL of Binding Buffer 1 and 500 μL of Binding Buffer 2 were added to the sample, respectively. The samples were then centrifuged at 3000 × g for 10 min at room temperature to remove the precipitate. Following protocols (DNA purification, elution, and collection) were identical to those for the sponge solution samples. All the eDNA samples were stored at −20°C until the first polymerase chain reaction (PCR) step in the eDNA metabarcoding.

### Quantitative fish environmental DNA metabarcoding

In this study, we used the quantitative eDNA metabarcoding (qMiSeq, Ushio et al. 2019) method to identify the fish species composition and quantify the fish eDNA concentrations simultaneously. In general, sequence reads obtained from high-throughput sequencing do not necessarily correspond to eDNA concentrations owing to the differences in PCR efficiency (primer bias, PCR inhibition, etc.). Applying the qMiSeq method, sequence reads can be converted into DNA concentrations (DNA copies/template) using sample-specific standard regression lines.

Amplicon libraries were prepared according to the following protocol (detailed information is provided in Supplementary Information S1). Fish eDNA from each sample was amplified using MiFish primers (MiFish-U, -O, and -L; Miya et al. 2020). An internal standard DNA mix (5, 25, and 50 copies/template) was added to each PCR mix. The first PCR was repeated eight times for each sample. The replicated samples were pooled as a single first PCR product for use in subsequent steps. In the first PCR step, PCR-negative controls (eight replicates) were included. In the second PCR, 8 bp dual indices were added to the purified first PCR amplicons. The second PCR amplicons were pooled, and the target library size (approximately 370 bp) of the second PCR was collected using E-Gel SizeSelect agarose gels (Thermo Fisher Scientific). The purified library was sequenced using the iSeq i1 Reagent and the iSeq 100 platform with 30% PhiX control (Illumina, San Diego, CA, USA). The sequencing dataset output from the iSeq platform was subjected to a bioinformatic analysis. After the analysis, sequence reads of each fish species and the standard DNA mix in each sample were obtained and subjected to a qMiSeq analysis. To obtain sample-specific regression lines, a regression analysis was performed using sequence reads and the DNA concentration of the internal standard DNA mix. The slopes and R^2^ values for each regression line are listed in Supplementary Tables S1–3. The R^2^ values of all the regression lines are higher than 0.98. In this study, PCR inhibition was not apparent in any of the water or passive DNA samples.

### Statistical analysis

In this study, the differences in the number of detected species between the two methods (water and passive sampling) were assessed in each experiment. In each experiment, to examine the differences in diurnal conditions, the number of species in water samples was separated into three categories: water (all), water (day), and water (night). Water (day) contained daytime samples (0–6 h and 20–24 h), water (night) contained nighttime samples (8 h–18 h), and water (all) contained all the water samples (day and night). The Kruskal–Wallis test was conducted to examine the differences in the number of species among all the survey protocols (three conditions of water samples and passive sampling methods) in each experiment. Steel–Dwass multiple comparisons were then performed in the experiments with significant differences.

Using the fish eDNA concentration dataset, non-metric multidimensional scaling (nMDS) in 1000 separate runs was performed to visualize the differences in fish community structure among the sampling methods. The stress value was used as an indicator of the nMDS fitting. The threshold stress values in the nMDS were less than 0.2, and the lowest value was selected. In addition, permutational multivariate analysis of variance (PERMANOVA) was used to examine the statistical differences in fish community structure among the sampling methods, and permutational analysis of multivariate dispersions (PERMDISP) was used to assess the variation in species composition among the sampling methods. The Bray–Curtis dissimilarity of fish community structure between samples in each experiment was calculated. The β-diversity plots of each sample were visualized in each experiment. In each experiment, fish species that affected the differences in community structure between sampling methods were estimated using the “envfit” function. In this study, the threshold p-value was < 0.01 for selecting species that had higher effects on the community structure. All the statistical analyses were performed using R ver. 4.02. For nMDS, PERMANOVA, and PERMDISP, the “metaMDS”, “adonis”, and “betadisper” functions of the vegan package were used.

## Results

In this study, 8,296,115 reads were obtained from 105 water samples and 41 passive samples, and 6,903,093 reads remained after preprocessing (Supplementary Table S4). The number of species detected in Experiment 1 (July 2021) ranged from 15 to 22 for the water samples (n = 39) and 7–14 (M-filter, n = 3) and 15–22 (M-solution, n = 8) for the passive samples (Fig. 2). Because of the loss of one of the M-solution samples in Experiment 1 (M-solution 3-3) from the equipment, the total number of M-solution samples in Experiment 1 was eight (Supplementary Table S1). The number of species detected in Experiment 2 (October 2021) ranged from 15 to 23 for the water samples (n = 39) and from 19 to 25 for the M-solution samples (n = 18). In Experiment 3 (May 2022), the number of species ranged from 18 to 24 for the water samples (n = 27), and 21–23 (M-solution, n = 3), 21–23 (S-solution, n = 3), 16–19 (S-syringe, n = 3), and 19 (S-syringe-NP, n = 3) for the passive samples (Fig. 2). Steel–Dwass multiple comparisons were performed because significant differences were verified by the results of the Kruskal–Wallis test in all the experiments (Supplementary Table S5). However, the results of the Steel–Dwass test verified significant differences only in Experiment 1 (Fig. 2). In that experiment, significant differences were observed between the number of species detected from M-filter and that of water (all), water (day), water (night) and between water (day) and water (night), respectively (*p* < 0.05). In Experiment 2, the number of species detected in the M-solution samples tended to be larger than that in water (all) and water (day), although the differences were not statistically significant (*p* = 0.055 and 0.072, respectively). In Experiment 3, there were no statistically significant differences in number of species among the sample types (Supplementary Table S5).

**Fig 2.**
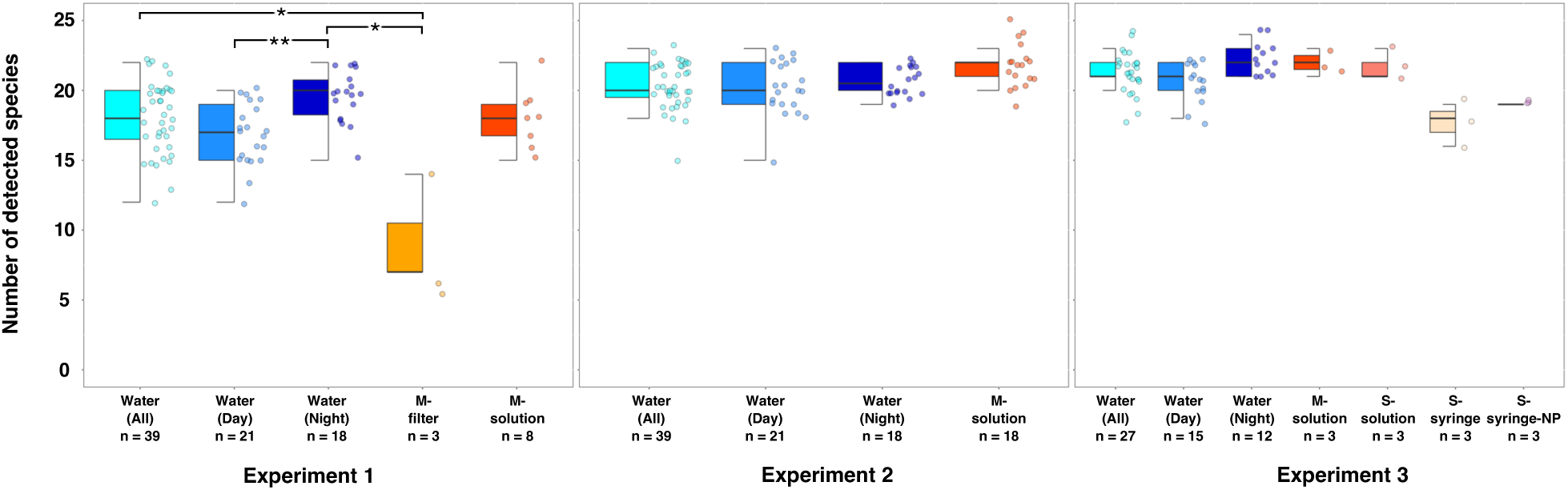
Boxplots in the number of detected species in each experiment. Asterisks indicate the significant differences among sample categories (**: *p* < 0.01, *: *p* < 0.05)

The cumulative fish diversity was higher for water sampling accumulated between 0 and 24 h (3 samples × 13 times) than for passive sampling methods in all the experiments (Fig. 3).

**Fig 3.**
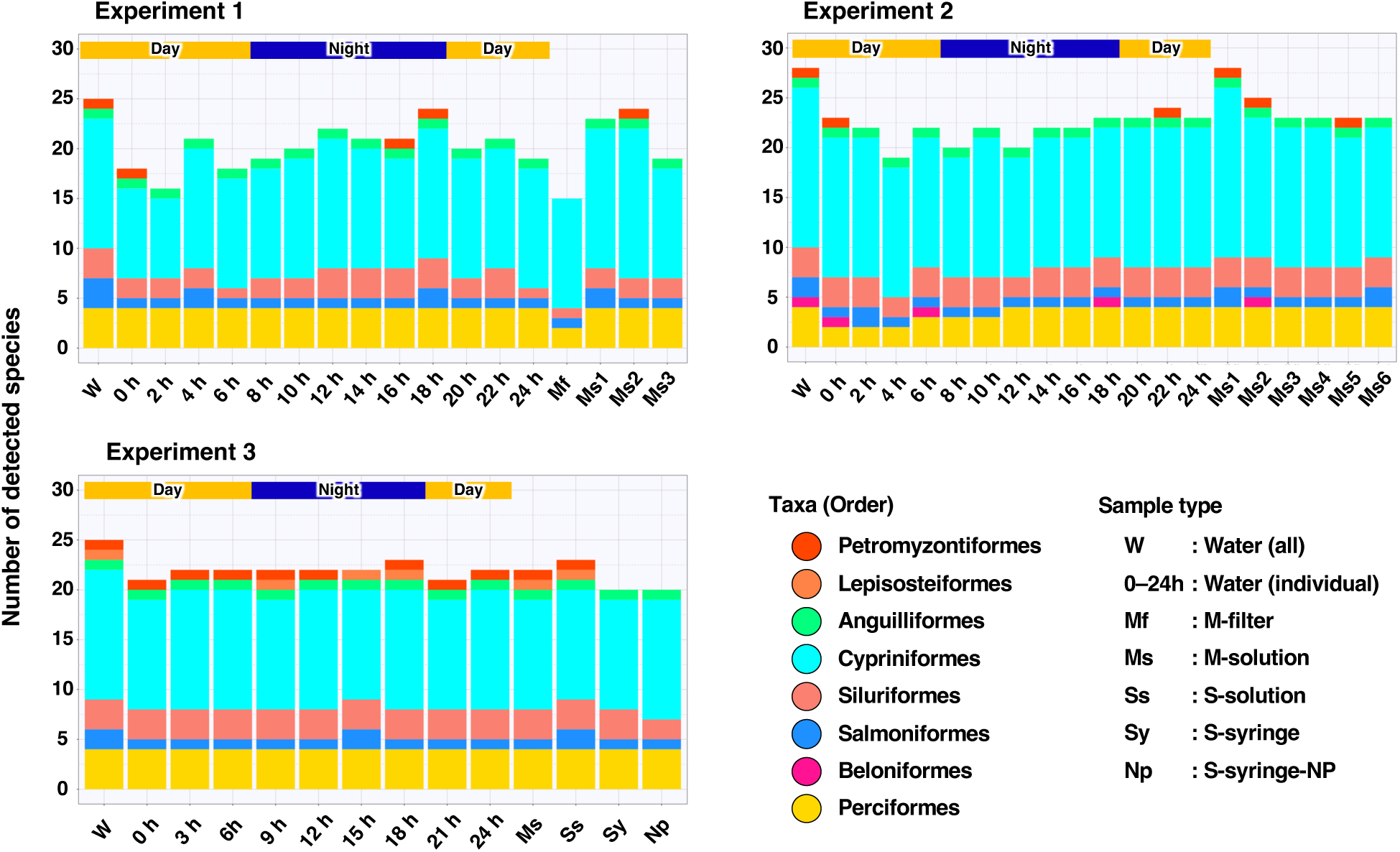
Cumulative number of species detected for each collection timing and DNA extraction method

Specifically, the cumulative numbers of fish species in Experiment 1 were 25, 15, 23, 24, and 19 in the water; M-filter; and M-solution 1, 2, and 3 samples, respectively. The cumulative number of fish species in Experiment 2 was 28 and 31 for the water and M-solution samples, respectively. In Experiment 3, the cumulative numbers of fish species in the water, M-solution, S-solution, S-syringe, and S-syringe-NP samples were 26, 23, 24, 21, and 21, respectively. Passive sampling methods also showed a high cumulative fish diversity similar to that for water. Herein, the M-solution in Experiment 2 showed the highest diversity (Supplementary Tables S1–3).

The fish species composition was substantially similar in all the experiments (Fig. 4). The number of species common between the water (day) or water (night) samples and the passive samples (M-solution, S-solution, S-syringe, or S-syringe-NP) was significantly larger than that for the other combinations in all the experiments. However, the common fish species in the two water samples (nine species) were not detected in the M-filter samples in Experiment 1 (Fig. 4).

**Fig 4.**
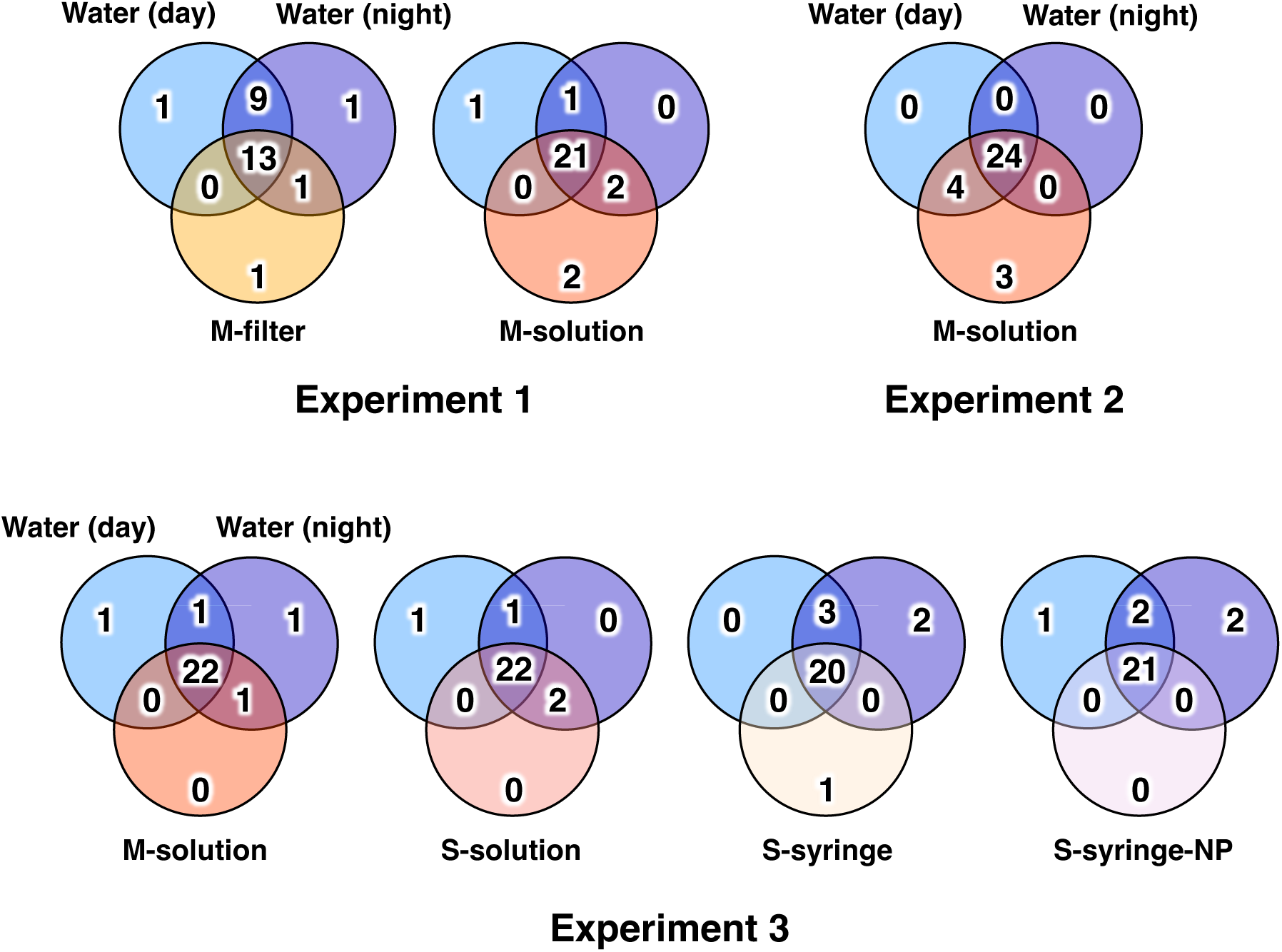
Venn diagrams showing the number of fish species belonging to each sampling method that were detected by eDNA metabarcoding in each experiment. Detailed information on the fish species compositions in each experiment is presented in Supplementary Table S6

Meanwhile, the comparison with the M-solution showed a decrease in the number of species detected only in the two water samples and an increase in the number of species common to the three methods compared with the M-filter. The cumulative fish species composition detected by each sampling method in Experiments 1–3 is listed in Supplementary Table S6. In Experiment 1, *Oncorhynchus masou* subsp. was specifically detected in water (day), and *Pseudogobio agathonectris* was detected in the passive samples (M-filter and M-solution). In Experiment 2, three fish species (*Acheilognathus rhombeus*, *Hemibarbus* sp., and *Salvelinus leucomaenis* subsp.) were detected only in the M-solution samples. The species detected only in the two types of water samples were not observed. In Experiment 3, *Tanakia limbata* was detected only in the water (day) samples, and *Hemibarbus* sp. was detected only in the S-syringe samples. In Experiment 3, the number of common species between the two types of water samples and the four passive sampling methods were similar. The fish species detected by eDNA metabarcoding have generally been known to inhabit the study sites in previous capturing surveys (River Environment Database System, 2018). However, the tropical garfish *Atractosteus tropicus* (listed as an alien species in Japan) was newly detected with high DNA concentrations in the water (night), M-solution, and S-solution samples.

In this study, an nMDS plot was used to assess the differences in community structure considering the eDNA concentration of each fish species (Fig. 5). The fish community structure in Experiment 1 differed significantly among the sample types (PERMANOVA: *p* < 0.01; PERMDISP: *p* = 0.441). In particular, the M-filter samples showed a higher variation than the other sample types. Although the number of species in the M-solution samples was similar to that in the two water samples, a few differences were observed in the community structure. Eight fish species affected the differences in community structure. Two of these species (*Hemibarbus longirostris* and *Tachysurus nudiceps*) were not detected in the M-filter samples. Other species were detected in all the sample types. However, their DNA concentrations were significantly higher in the water samples than in the passive samples (Supplementary Table S1). In Experiment 2, the fish community structure in water (night) was included in that of water (day), and the M-solution fish community was plotted outside the two types of water samples. However, differences in community structure among the three types could not be assessed precisely (PERMANOVA, *p* < 0.01; PERMDISP, *p* < 0.05). Differences in community structure between the water (all) and M-solution samples could not be assessed (PERMANOVA, *p* < 0.01; PERMDISP, *p* < 0.05). Fourteen fish species affected the differences in community structure in Experiment 2. *Hemibarbus* sp. Was detected only in the M-solution samples. Other species were detected in all the sample types. The trend of the DNA concentration of these species was similar to that observed in Experiment 1 (Supplementary Table S2). In Experiment 3, the fish community structure differed marginally between the water (day) and nighttime water (night) samples. In the passive samples, the community structure differed between solution types (M-solution and S-solution) and syringe types (S-syringe and S-syringe-NP). In Experiment 3, the fish community structure differed significantly among the sample types (PERMANOVA: *p* < 0.01, PERMDISP: *p* = 0.11). Thirteen fish species affected the community structure in Experiment 3. All the species were detected in all the sample types, except for Misgurnus anguillicaudatus in the S-syringe samples. The trend of the DNA concentration of these species was similar to that observed in Experiments 1 and 2 (Supplementary Table S3).

**Fig 5.**
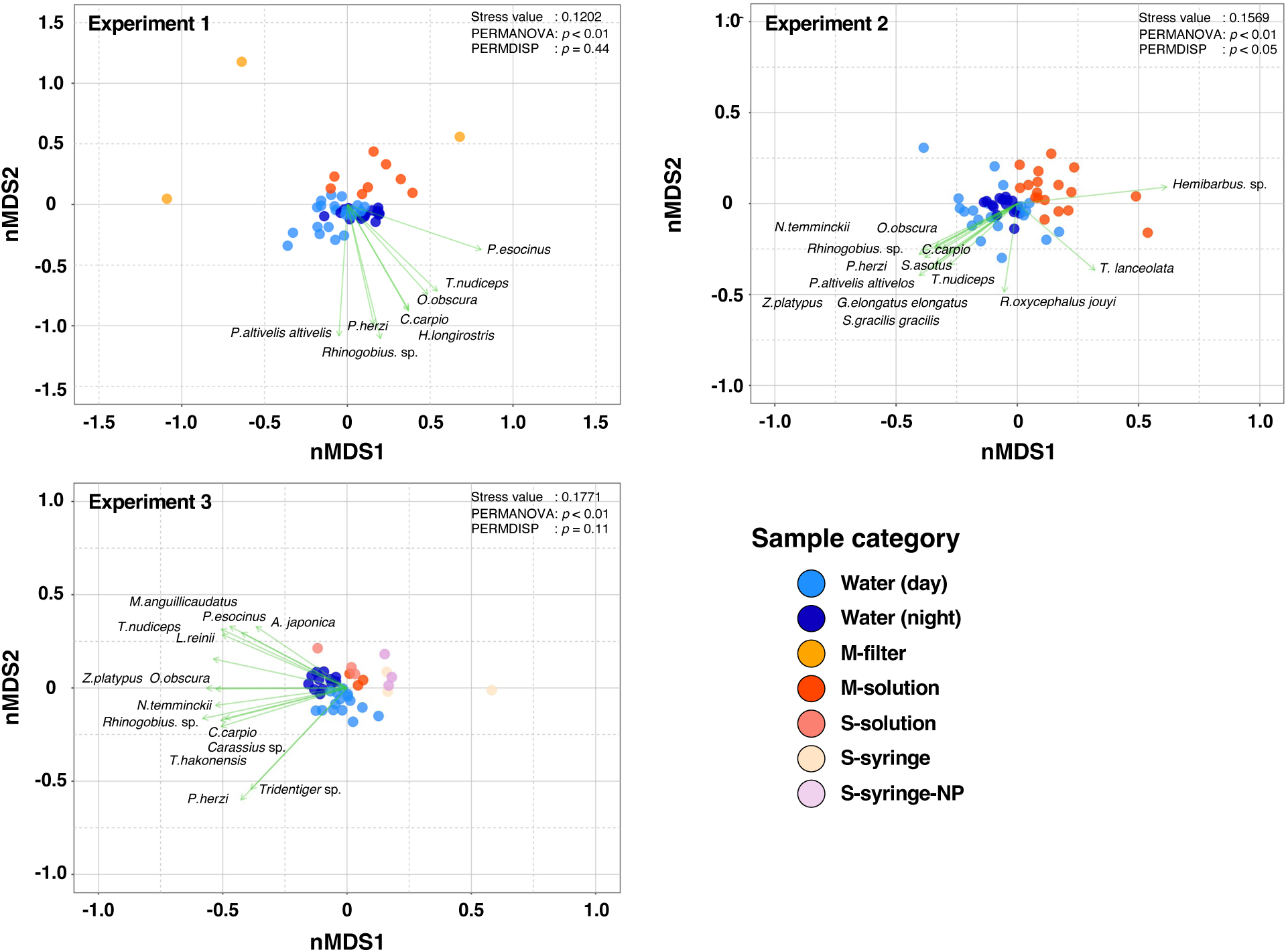
nMDS plots for each experiment. The green arrows represent the fish responsible for causing a significant difference among sampling methods.

## Discussion

In this study, we assessed the effectiveness of passive eDNA sampling methods compared with water sampling in a riverine environment. In addition, in the passive sampling method, several DNA extraction methods were examined to determine the optimal method for eDNA recovery from sponges. Our results showed that the passive sampling method using sponge skeletons had a similar or larger number of fish species than water samples. The results of the present study support previous studies showing similar trends (Jaunen et al. 2024, Zhan et al. 2024, Mlinarec et al. 2025). However, in Experiment 1, the number of species in the M-filter sample was significantly less than that for the other methods (water and other passive samples), and the variation in fish community composition was larger than that in the other methods. Meanwhile, because no PCR inhibition was observed in any of the DNA samples, other factors may have affected the reduction in the number of species in the M-filter samples. These results indicate that the DNA recovery rate of M-filter was less than that of the other methods. Water samples pressed from sponges contained large amounts of material in addition to eDNA (e.g., sand, mud, and biofilm). The DNA in pressed solutions could be captured by these materials. This could be assumed straightforwardly from previous studies using soil particles and biofilms as capturing materials for passive sampling (Kirtane et al., 2020, Rivera et al., 2021). The method of DNA extraction from the GF/F filter used in the M-filter may not have efficiently recovered the eDNA captured from these materials. In contrast, the DC method applied to the passive samplers in this study indicated that the efficiency of DNA recovery was improved by direct DNA extraction from pressed solutions or by direct immersion of sponge skeletons in the elution mixture. The M-solution in the passive sampling method showed similar fish diversity in all the experiments (in terms of both number of species and cumulative number of species) similarly as in the water samples. Furthermore, in Experiment 2, fish species detected only in M-solution were observed. The cumulative number of samples in the passive sampling methods was larger than that for the water samples (Fig. 4). These results show that passive samples (sponge skeletons) with 24 h immersion displayed detection performance equal to that for 24 h water samples extracted every 2 h. The smaller number of species observed for the other passive sampling methods (S-solution, S-syringe, and S-syringe-NP) was marginally less than that in the water samples. However, statistically significant differences were not observed. The high cumulative fish diversity for the S-solution and the presence of specific species in the S-syringe may also indicate the effectiveness of each method (Figs. 3 and 4). However, it could have been difficult to distinguish the passive sampling method in Experiment 3 from the other methods because of the absence of replicate samples (triplicates in each sample). Therefore, the effectiveness of these methods should be evaluated meticulously in future studies.

In Experiments 1 and 3, the number of species detected in the water samples at nighttime tended to be larger than that for the day time (Fig. 3). The cumulative number of species in the water samples in Experiments 1 and 2 varied considerably with collection timing (Fig. 4). In contrast, the number of detected and cumulative species in the passive samples, except for the M-filter sample, tended to be more homogeneous than those in the water samples in all the experiments. These differences could be owing to dissimilarities in the DNA information of freshwater fish between the water and passive samples. In general, the eDNA information from water samples contains snapshot information that reflects the collection timing (Spear et al. 2015, Bowers et al., 2021; Jensen et al., 2022). Therefore, eDNA removed from rare species or species with an incidental appearance can be rapidly diluted and dispersed and could elude detection via instantaneous water sampling (Kirtane et al., 2020). In contrast, passive sampling methods could accumulate more eDNA by filtering a large water volume through long-term immersion in water. Jeunen et al. (2022) indicated that more porous passive filter materials could achieve higher detection rates through the adsorption and retention of low-abundance DNA and DNA bound to particulate matter. The differences in temporal resolution between sampling methods could be reflected in these variations and differences in fish species diversity.

Experiments 1 and 3 showed differences in fish community structure between the methods in the nMDS plot and statistical analyses. In all the experiments, certain fish species affected the differences in community structure among the sample types. In Experiment 1, this was reasonable because the fish diversity and DNA concentrations of each fish species in the M-filter samples were significantly lower than those in the other methods. However, in the other experiments, the fish diversity and species composition differed among the sampling methods (Figs. 3 and 4). Therefore, the variations in community structure among the different methods could be attributed to the differences in the eDNA concentration of each fish species. There were differences in fish DNA concentrations among the sampling methods. Those of the common fish species (*Zacco platypus*, *Nipponocypris temminckii*, and *Pungtungia herzi*) observed at the survey site showed large differences between the water and passive samples in all the experiments (Supplementary Tables 1– 3). The fish DNA concentration of each sample was standardized by adding the standard DNA mix. Therefore, these differences indicated that the eDNA concentration of fish in the passive samples was lower than that in the water samples. Passive sampling methods store aquatic eDNA by capturing and accumulating collection material. Cananzi et al. (2025) showed that passive samplers contain eDNA of diverse taxa and particle sizes and are more efficient for detecting micro-invertebrates and amphibians than water samples using a multi-marker approach. Similarly, in our study, large amounts of eDNA from non-fish organisms (e.g., aquatic plants, insects, and microorganisms) may have been captured by sponge skeletons during the long-term immersion. As a result, fish eDNA was relatively reduced in the passive samples. This generated differences in the proportion of DNA concentration between the water samples and passive samples. The differences in eDNA concentrations may also reflect the differences in DNA extraction efficiency between the sponge skeletons and GF/F filters. Although the optimization of DNA extraction procedures using GF/F filters has been examined in previous studies (Minamoto et al. 2021), the optimization of sponge skeletons may still be insufficient. Previous studies have also shown that the efficiency of DNA recovery from passive samples varies significantly depending on the sampler material and the immersion method (Verdier et al. 2022, Saltonstall et al. 2024, Stevens et al. 2024). Therefore, future improvements in the methodology would enhance the efficacy of the passive sampling method using sponge skeletons.

To conclude, we showed that passive eDNA sampling using sponge skeletons is one of the most efficient methods for monitoring freshwater fish in riverine environments. Passive sampling methods enable more cost-effective and efficient eDNA monitoring than the conventional filtering methods. Passive sampling methods also do not require time-consuming filtration or the preservation of large volumes of water. These methods minimize sampling and processing times in the field, accumulate eDNA through long-term immersion, and enable time-efficient biomonitoring. Our passive sampling method could detect similar or higher fish species diversity than water sampling because of the long-term and cumulative eDNA adsorption on sponge skeletons.

However, there are challenges associated with passive sampling methods. One of these is that the actual amount of water passing through the passive collection device during submersion is difficult to quantify (Zhang et al. 2024). Currently, this method may be unsuitable for estimating the eDNA concentration of target species or for quantitative assessments of species abundance and biomass. In this study, the eDNA concentration of each fish species was estimated using the qMiSeq method. However, the accuracy of DNA quantification could not be evaluated. This problem can be resolved by measuring the flow rate and estimating the water volume through the sponge skeleton. Furthermore, passive sampling methods can concentrate PCR inhibitors and eDNA through long-term immersion. Organic compounds (e.g., humic and fulvic acid) produced by the biological processes of phytoplankton and terrestrial detritus inputs are present in aquatic systems and can be the major cause of PCR inhibition (Hellings et al. 1999, Albers et al., 2013, Uchii et al. 2019). These substances may be adsorbed and concentrated on the sponge skeleton, as well as on eDNA. This would increase the probability of PCR inhibition. Further improvements in methodology, such as the effect of PCR inhibition, quantitative assessment, and immersion period, are required so that passive sampling methods using sponge skeletons can be an effective tool for eDNA surveys.

## Acknowledgments

We thank the students of the Akamatsu Laboratory at Yamaguchi University for their support in the field experiments. We also thank the Saba River Fisheries Cooperative and Yamaguchi River and National Highway Office, Chugoku Regional Development Bureau, Ministry of Land, Infrastructure, Transport, and Tourism for granting permission to conduct field surveys. This work was supported by a Grant-in-Aid for Challenging Research (Exploratory) from the Japan Society for the Promotion of Science (Grant ID: 22K18298).

## Data accessibility

The entire sequence data would be published in DRA (Accession number:) after the manuscript is accepted. Detailed information on the fish community structure and methodological descriptions of library preparation are available in the Supplementary Materials.

## References

Albers CN, Jensen A, Bælum J, Jacobsen CS (2013) Inhibition of DNA polymerases used in Q-PCR by structurally different soil-derived humic substances. Geomicrobiol J 30:675–681. 10.1080/01490451.2012.758193

Bessey C, Jarman SN, Simpson T, Miller H, Stewart T, Keesing JK, Berry O (2021) Passive eDNA collection enhances aquatic biodiversity analysis. Commun Biol 4:236. 10.1038/s42003-021-01760-8

Bowers HA, Pochon X, von Ammon U, Gemmell N, Stanton JAL, Jeunen GJ, Sherman CDH, Zaiko A (2021) Towards the optimization of eDNA/eRNA sampling Technologies for Marine Biosecurity Surveillance. Water 13(8):13081113. 10.3390/w13081113

Cai W, Harper LR, Neave EF, Shum P, Craggs J, Arias MB, Riesgo A, Mariani S (2022) Environmental DNA persistence and fish detection in captive sponges. Mol Ecol Resour 22:2956–2966. 10.1111/1755-0998.13677

Cananzi G, Tatini I, Li T, Montagna M, Serra V, Petroni G (2025) Active or passive? A multi-marker approach to compare active and passive eDNA sampling in riverine environments. Sci Total Environ 974:179247. 10.1016/j.scitotenv.2025.179247

Hänfling B, Handley LL, Read DS, Hahn C, Li J, Nichols P, Blackman RC, Oliver A, Winfield IJ (2016) Environmental DNA metabarcoding of lake fish communities reflects long-term data from established survey methods. Mol Ecol 25:3101–3119. 10.1111/mec.13660

Hellings L, Dehairs F, Tackx M, Keppens E, Baeyens W (1999) Origin and fate of organic carbon in the freshwater part of the Scheldt Estuary as traced by stable carbon isotope composition. Biogeochemistry 47:167–186. 10.1023/A:1006143827118

Jensen MR, Sigsgaard EE, de Ávila M P, Agersnap S, Brenner-Larsen W, Sengupta ME, Xing Y, Krag MA, Knudsen SW, Carl H, Møller PR, Thomsen PF (2022) Short-term temporal variation of coastal marine eDNA. Environ DNA 4:747–762. 10.1002/edn3.285

Jeunen GJ, Knapp M, Spencer HG, Taylor HR, Lamare MD, Stat M, Bunce M, Gemmell NJ (2019) Species-level biodiversity assessment using marine environmental DNA metabarcoding requires protocol optimization and standardization. Ecol Evol 9:1323–1335. 10.1002/ece3.4843

Jeunen GJ, Mills S, Mariani S, Treece J, Ferreira S, Stanton JAL, Duŕan-Vinet B, Duffya GA, Gemmell NJ, Lamare M (2024) Streamlining large-scale oceanic biomonitoring using passive eDNA samplers integrated into vessel’s continuous pump underway seawater systems. Sci Total Environ 946:174354. 10.1016/j.scitotenv.2024.174354

Jeunen GJ, von Ammon U, Cross H, Ferreira S, Lamare M, Day R, Treece J, Pochon X, Zaiko A, Gemmell NJ, Stanton JA (2022) Moving environmental DNA (eDNA) technologies frombenchtop to the field using passive sampling and PDQeXextraction. Environ DNA 4:1420– 1433. 10.1002/edn3.356

Kirtane A, Atkinson JD, Sassoubre L (2020). Design and validation of passive environmental DNA samplers using granular activated carbon and montmorillonite clay. Environ Sci Technol 54:11961–11970. 10.1021/acs.est.0c01863

Lynggaard C, Bertelsen MF, Jensen CV, Johnson MS, Frøslev TG, Olsen MT, Bohmann K (2022) Airborne environmental DNA for terrestrial vertebrate community monitoring. Curr Biol 32:701– 707. 10.1016/j.cub.2021.12.014

Mariani S, Baillie C, Colosimo G, Riesgo A (2019) Sponges as natural environmental DNA samplers. Curr Biol 29(11):R401–R402. 10.1016/j.cub.2019.04.031

Minamoto T, Miya M, Sado T, Seino S, Doi H, Kondoh M, Nakamura K, Takahara T, Yamamoto S, Yamanaka H, Araki H, Iwasaki W, Kasai A, Masuda R, Uchii K (2021) An illustrated manual for environmental DNA research: Water sampling guidelines and experimental protocols. Environ DNA 3:8–13. 10.1002/edn3.121

Ministry of Land, Infrastructure, Transport and Tourism, Japan. (2018) Chugoku District in River Environment Database system. https://www.nilim.go.jp/lab/fbg/ksnkankyo/dl_87_index.html (referred in October 10, 2022)

Miya M, Gotoh RO, Sado T (2020) MiFish metabarcoding: A high - throughput approach for simultaneous detection of multiple fish species from environmental DNA and other samples. Fish Sci 86:939–970. 10.1007/s12562-020-01461-x

Mlinarec J, Svetličić I, Kresonja M, Rubinić M, Megyery T, Kaliger I, Mihaljević D (2025) Temporal and spatial dynamics of the lotic fish communities: A comparison of coffee filter-based Passive eDNA collection versus active eDNA filtering. Environ DNA 7. 10.1002/edn3.70065o

Nakagawa H, Yamamoto S, Sato Y, Sado T, Minamoto T, Miya M (2018) Comparing local- and regional-scale estimations of the diversity of stream fish using eDNA metabarcoding and conventional observation methods. Freshw Biol 63:569–580. 10.1111/fwb.13094

Oka SI, Doi H, Miyamoto K, Hanahara N, Sado T, Miya M (2021) Environmental DNA metabarcoding for biodiversity monitoring of a highly diverse tropical fish community in a coral reef lagoon: Estimation of species richness and detection of habitat segregation. Environ DNA 3:55–69. 10.1002/edn3.132

Rivera SF, Rimet F, Vasselon V, Vautier M, Domaizon I, Bouchez A (2021). Fish eDNA metabarcoding from aquatic biofilm samples: Methodological aspects. Mol Ecol Res 22:1440– 1453. 10.1111/1755-0998.13568

Saltonstall K, Delgado J, Vargas M, Collin R (2024) Are passive collectors effective samplers of microbes in natural aquatic systems? Front. Freshw Sci 2:1460713. 10.3389/ffwsc.2024.1460713

Shaw JL, Clarke LJ, Wedderburn SD, Barnes TC, Weyrich LS, Cooper A (2016) Comparison of environmental DNA metabarcoding and conventional fish survey methods in a river system. Biol Conserv 197:131–138. 10.1016/j.biocon.2016.03.010

Stevens ER, Hyde J, Beesley LS, Gwinn DC, Thompson S, Morris L, Wilson PR, Gleeson DB (2024) Fishy Business—Assessing the efficacy of active and passive eDNA to describe the fish assemblage of a river in Southwestern Western Australia to support effective monitoring. Environ DNA 6:6. 10.1002/edn3.70040

Tsuji S, Inui R, Nakao R, Miyazono S, Saito M, Kono T, Akamatsu Y (2022) Quantitative environmental DNA metabarcoding shows high potential as a novel approach to quantitatively assess fish community. Sci Rep 12:21524. 10.1038/s41598-022-25274-3

Tsuji S, Takahara T, Doi H, Shibata N, Yamanaka H (2019) The detection of aquatic macroorganisms using environmental DNA analysis—A review of methods for collection, extraction, and detection. Environ DNA 1:99–108. 10.1002/edn3.21

Turner CR, Uy KL, Everhart RC (2015) Fish environmental DNA is more concentrated in aquatic sediments than surface water. Biol Conserv 183:93–102. 10.1016/j.biocon.2014.11.017

Turon M, Angulo-Preckler C, Antich A, Præbel K, Wangensteen OS (2020). More than expected from old sponge samples: A natural sampler DNA metabarcoding assessment of marine fish diversity in Nha Trang Bay (Vietnam). Front Mar Sci 7:1–14. 10.3389/fmars.2020.605148

Uchii K, Doi H, Okahashi T, Katano I, Yamanaka H, Sakata MK, Minamoto T (2019) Comparison of inhibition resistance among PCR reagents fordetection and quantification of environmental DNA. Environ DNA 1:359–367. 10.1002/edn3.37

Ushio M (2019) Use of a filter cartridge combined with intra-cartridge bead-beating improves detection of microbial DNA from water samples. Methods Ecol Evol 10:1142–1156. 10.1111/2041-210X.13204

Ushio M, Murakami H, Masuda R, Sado T, Miya M, Sakurai S, Yamanaka H, Minamoto T, Kondoh M (2018) Quantitative monitoring of multispecies fish environmental DNA using high-throughput sequencing. Metabarcoding Metagenom 2:e23297. 10.3897/mbmg.2.23297

Verdier H, Konecny-Dupre L, Marquette C, Reveron H, Tadier S, Grémillard L, Barthès A, Datry T, Bouchez A, Lefébure T (2022) Passive sampling of environmental DNA in aquatic environments using 3D-printed hydroxyapatite samplers. Mol Ecol Resour 22:2158–2170. 10.1111/1755-0998.13604

Yamamoto S, Masuda M, Sato Y, Sado T, Araki H, Kondoh M, Minamoto T, Miya M (2017) Environmental DNA metabarcoding reveals local fish communities in a species-rich coastal sea. Sci Rep 7:40368.

Yan Z, Luo Y, Chen X, Yang L, Yao M (2024) Angling and trolling for eDNA: A novel and effective approach for passive eDNA capture in natural waters. Environ Int 194:109175. 10.1016/j.envint.2024.109175

Zhang L, Zhou W, Jiao M, Xie T, Xie M, Li H, Suo A, Yue W, Ding D, He W (2024) Use of passive sampling in environmental DNA metabarcoding technology: Monitoring of fish diversity in the Jiangmen coastal waters. Sci Total Environ 908:168298. 10.1016/j.scitotenv.2023.168298

Zinger L, Bonin A, Alsos IG et al. (2019) DNA metabarcoding—Need for robust experimental designs to draw sound ecological conclusions. Mol Ecol 28:1857–1862.

