## Supplementary Information for "Efficacy of natural marine sponges as a passive environmental DNA sampler for freshwater fish diversity monitoring"

**Title:**

**Evaluation of the efficacy of natural sponges as a passive environmental DNA (eDNA) sampler for freshwater biodiversity monitoring**

Author’s Name: **Ryohei Nakao^1^, Manami Inaba^1^, Seiji Miyazono^1^, Minoru Saito^1,2^, Keita Maruyama^1,4^, Fumiko Imamura^3^, Yoshihisa Akamatsu^1^**

Institution:

^1^ Graduate School of Science and Technology for Innovation, Yamaguchi University, Yamaguchi University, 2‑16‑1, Tokiwadai, Ube, Yamaguchi, Japan

^2^ Japan International Research Center for Agricultural Sciences, 1-1 Ohwashi, Tsukuba, Ibaraki, Japan

^3^ Research and Development Center, Nippon Koei Co., Ltd, 2304, Inarihara, Tsukuba, Ibaraki, Japan

^4^ Department of Ocean Sciences, Tokyo University of Marine Science and Technology, 4-5-7 Konan, Minato-ku, Tokyo 108-8477, Japan

**Supplementary Information S1: Further methodological details of tha quantitative environmental DNA metabarcoding**

***Library preparation and metabarcoding using qMiSeq method***

The amplicon library was prepared according to the following protocols. In the first PCR, the total reaction volume was 12 μL, containing 6.0μL of 2× KAPA HS ready Mix (Nippon Genetics, Tokyo, Japan), 0.72 μL of each MiFish-U primers (5µM), 0.36 µL of two sub-primers (5µM MiFish-O and L) (Miya et al. 2020), 1.12 µL of ultrapure water, 1.0 µL of standard DNA mix (containing 5 copies, 25 copies, and 50 copies/µL) and 1.0 μL of template DNA. The thermocycling conditions were 94℃ for 2 min, 35 cycles of 98℃ for 10 s, 65℃ for 30 s, 68℃ for 30 s, and 68℃ for 5 min. The first PCR was repeated eight times for each sample, and the replicated samples were pooled as a single first PCR product for use in the subsequent step. The pooled first PCR products were purified using GeneRead™ Size Selection Kit (QIAGEN, Hilden, Germany) according to the manufacturer’s instructions. The DNA concentrations of purified first PCR products were measured using a Qubit dsDNA HS assay kit and a Qubit 3.0 fluorometer (Thermo Fisher Scientific, Waltham, MA, USA). All purified first PCR products were diluted to 0.1 ng/μL with ultrapure water, and the diluted samples were used as templates for the following second PCR. In the first PCR step, the PCR negative controls (eight replicates) were included in each experiment.

The second PCR was performed to add adapter sequences with 8-bp dual indices. The total reaction volume was 12 μL, containing 6.0 μL of 2× KAPA HiFi HS ReadyMix, 2.0 μL of forward and reverse primer (1.8 μM), 1.0 μL of purified first PCR product, and 1.0 μL of ultrapure water. The thermocycling conditions were 95℃ for 3 min, 12 cycles of 98℃ for 20 s, 72℃ for 15 s, and 72℃ for 5 min. Indexed PCR products were pooled in the equivalent volume, and 25 μL of the pooled libraries were loaded on a 2% E-Gel SizeSelect agarose gels (Thermo Fisher Scientific), and a target library size (ca. 370 bp) was purified. The quality of the amplicon library was checked using an Agilent 4150 TapeStation (Agilent Technologies Inc., Santa Clara, CA, USA), and the DNA concentrations of the amplicon library were measured using Qubit dsDNA HS assay Kit using a Qubit 3.0 fluorometer.

Amplicon library was sequenced using iSeq i1 Reagent and iSeq 100 platforms (Illumina, San Diego, CA, USA). 20 µL of 30 pM amplicon library was spiked with approximately 30% PhiX control (PhiX Control Kit v3, Illumina) before sequencing runs. Subsequently, the sequencing dataset output from iSeq was subjected to pre-processing and taxonomic assignments. All sequence data are registered in the DNA Data Bank of Japan (DDBJ) Sequence Read Archive (DRA, Accession number: DRAXXXXX).

***Denoising and taxonomic assignment***

We used the USEARCH v11.0667 for all data pre-processing activities and taxonomic assignment of the sequence dataset obtained from the iSeq. First, pair-end reads (R1 and R2 reads) generated from iSeq was assembled using the “fastq_mergepairs” command (overlapped reads are not written). In the process, the low-quality tail reads with a cut-off threshold at a Phred score of 2, and the paired reads with too many mismatches (> 5 positions) in the aligned regions were discarded. Secondly, the primer sequences were removed from the merged reads using the “fastx_truncate” command. Afterward, read quality filtering was performed using the “fastq_filter” command with thresholds of max expected error > 1.0 and > 50 bp read length. The pre-processed reads were dereplicated using the “fastx_uniques” command, and the chimeric reads and less than 10 reads were removed from all samples as the potential sequence errors. Finally, an error-correction of amplicon reads, which checks and discards the PCR errors and chimeric reads, was performed using the “unoise3” command in the unoise3 algorithm32. Before the taxonomic assignment, the processed reads from the above steps were subjected to sequence similarity search using the “usearch_global” command against reference databases of fish species that had been established previously (MiFish local database v37). The sequence similarity and cut off E-value were 98.5 % and 10-5, respectively. If there was only one species with ≧ 98.5 % similarity, the sequence was assigned to the top-hit species. Conversely, sequences assigned to two or more species in the ≧ 98.5 % similarity were merged as species complex and listed in the synonym group. Generally, the species complexes were assigned to the genus level (e.g., Asian crucian carp Carassius spp.). Species that were unlikely to inhabit Japan and marine fishes were excluded from the candidate list of species complexes. Finally, sequence reads of each fish species were arranged into the matrix, with the rows and columns representing the number of sites and fish species (or genus), respectively.

**References**

Edgar RC (2010) Search and clustering orders of magnitude faster than BLAST. Bioinformatics 26:2460–2461. doi: 10.1093/bioinformatics/btq461

Hatama T, Hamano T, Saito M (2018) Distribution patterns of freshwater fishes and decapod crustaceans at biogeographical, river system, and segment scales in Yamaguchi Prefecture, western Japan. Bull. Biogeoger. Soc. Japan 72:141–199. (in Japanese)

Miya M, Sato Y, Fukunaga T, Sado T, Poulsen JY, Sato K, Minamoto T, Yamamoto S, Yamanaka H, Araki H, Kondoh M, Iwasaki W (2015) MiFish, a set of universal PCR primers for metabarcoding environmental DNA from fishes: detection of more than 230 subtropical marine species. Royal Society Open Science 2 (7). doi: 10.1098/rsos.150088

Oka S, Doi H, Miyamoto K, Hanahara N, Sado T, Miya M (2020) Environmental DNA metabarcoding for biodiversity monitoring of a highly diverse tropical fish community in a coral reef lagoon: Estimation of species richness and detection of habitat segregation. EnvironmentalDNA Early view. doi: 10.1002/edn3.132
